# A single vesicle fluorescence-bleaching assay for multi-parameter analysis of proteoliposomes by total internal reflection fluorescence microscopy

**DOI:** 10.1101/2022.04.01.486744

**Authors:** Sarina Veit, Laura Charlotte Paweletz, Sören S.-R. Bohr, Anant K. Menon, Nikos S Hatzakis, Thomas Günther Pomorski

## Abstract

Reconstitution of membrane proteins into model membranes is an essential approach for their functional analysis under chemically defined conditions. Established model-membrane systems used in ensemble average measurements are limited by sample heterogeneity and insufficient knowledge of lipid and protein content at the single vesicle level, which limits quantitative analysis of vesicle properties and prevents their correlation with protein activity. Here, we describe a versatile total internal reflection fluorescence microscopy-based bleaching protocol that permits parallel analyses of multiple parameters (physical size, tightness, unilamellarity, membrane protein content and orientation) of individual proteoliposomes prepared with fluorescently tagged membrane proteins and lipid markers. The approach makes use of commercially available fluorophores including the commonly used nitrobenzoxadiazole (NBD) dye and may be applied to deduce functional molecular characteristics of many types of reconstituted fluorescently tagged membrane proteins.

## Introduction

Reconstitution of purified proteins into model membranes is an essential approach for investigating membrane protein function under defined conditions outside the complex cellular environment (Rigaud and Lévy, 2003; Murray et al., 2014; Amati et al., 2020). Typically, proteins are reconstituted with phospholipids into large unilamellar vesicles (LUVs) with diameters on the order of 100 to 200 nm. These vesicles can be customized in a variety of ways, by using different lipid compositions to promote protein stability and/or control protein function, and including modified lipids such as pH-sensors, fluorophores or immobilization anchors to tailor the LUVs for a variety of applications. The bulk properties of such systems are readily determined by measuring lipid and protein content and sizing the vesicles by dynamic light scattering (DLS), nanoparticle tracking or electron microscopy. However, it is well established (Thomsen et al., 2019; Larsen et al., 2011; Lohse et al., 2008; Mathiasen et al., 2014; Guha et al., 2021; Kuyper et al., 2006; Thomsen et al., 2018) that individual vesicles within the bulk sample differ in size/curvature, permeability, lipid composition, protein content and orientation of reconstituted proteins. This high heterogeneity hampers quantitative analysis of vesicle properties (including stoichiometry of lipids and transmembrane proteins) and their correlation with protein activity. Other technical issues with commonly used ensemble measurements include potential leakiness of proteoliposomes and aggregation – these can skew the results and lead to their misinterpretation. To overcome these problems with ensemble assays, it is necessary to obtain information on a large and representative number of individual vesicles, for example, by using single vesicle microscopy. Previous work along these lines showed that the size distribution of the vesicles could be determined by correlating the fluorescence intensity of a lipid marker in single vesicles with DLS data (Hatzakis et al., 2009; Malle et al., 2021; Lohr et al., 2009; Olsson et al., 2015), and that tightness and lamellarity could be assessed by bleaching of the marker (Heider et al., 2011). Other studies used step-wise photobleaching assays to determine the copy number of fluorescent proteins per vesicle (Hummert et al., 2021). Here, we present a broadly applicable proteoliposome characterization assay at the single vesicle level that permits parallel analysis of multiple parameters (physical size, tightness, unilamellarity, membrane protein content and orientation) of individual proteoliposomes to provide a detailed picture of the reconstituted membrane system.

## Methods

### Materials

1-palmitoyl-2-oleoyl-*sn*-glycero-3-phosphocholine (POPC), 1-palmitoyl-2-oleoyl-*sn*-glycero-3-phospho-(1’-rac-glycerol) (sodium salt) (POPG), 1,2-distearoyl-*sn*-glycero-3-phosphoethanolamine-N-[biotinyl-(polyethylene glycol)-2000] (Biotinyl-PEG-DSPE) and *N*-(7-nitrobenz-2-oxa-1,3-diazol-4-yl)dioleoylphosphatidylethanolamine (*N*-NBD-DOPE) were obtained from Avanti Polar lipids Inc. (Birmingham, AL, USA). SNAP-Surface® Alexa Fluor® 647, SNAP-Surface® Alexa Fluor® 488 and SNAP-Cell® 647-SIR were obtained from New England Biolabs (Ipswich, MA, USA). Bio-Beads™ SM-2 Resin were obtained from Bio-Rad Laboratories Inc. (Hercules, CA, USA). The detergents n-Dodecyl-β-D-maltoside (DDM) and n-Octyl-β-D-Glucoside (OG) were obtained from GlyconBiochemicals GmbH (Luckenwalde, Germany). For surface coating, PLL(20)-g[3.5]-PEG(2) and PLL(20)-g[3.5]-PEG(2)/PEG(3.4)-biotin(20%) from SUSOS (Dübendorf, Switzerland) were used. The ionophores valinomycin and m-chlorophenylhydrazon (CCCP), the pH sensitive dye 9-amino-6-chloro-2-methoxyacridine (ACMA), and all other chemicals and regents were from Sigma-Aldrich (München, Germany), if not stated otherwise.

### Liposome preparation

POPC/POPG/Biotinyl-PEG-DSPE/*N*-NBD-DOPE (89.55/10/0.1/0.35 mol%) were mixed in chloroform and dried under vacuum (250 mbar) for minimum 3 h. Residual solvent was removed in high vacuum (20 mbar) for 30 min. The lipid film was resuspended in reconstitution buffer (20 mM MOPS-KOH, pH 7.0, 50 mM K_2_SO_4_) to a final concentration of 15 mg mL^−1^ by vortexing with a 5 mm glass bead for 10 min at 60°C followed by five freeze-thaw cycles (liquid nitrogen, 1 min; water bath at 60°C, 1.5 min). Vesicles were extruded through two nucleopore polycarbonate membranes with a pore size of 200 nm using a mini-extruder (Avanti Polar Lipids), yielding large unilamellar vesicles (LUVs).

### Protein purification

A 73 amino acid C-terminal truncated version of *Arabidopsis thaliana* auto-inhibited H^+^-ATPase isoform 2 (AHA2) containing a SNAP® tag in combination with StrepII and hexahistidine (6×His) affinity tags at the N-terminus of the protein was heterologously overexpressed in *Saccharomyces cerevisiae* strain RS-72 (Cid et al., 1987) and purified according to previously published protocols (Lanfermeijer et al., 1998). The purified protein (5 - 10 mg mL^−1^) in glycerol-containing buffer (50 mM MES-KOH, pH 6.5, 20 % (w/v) glycerol, 50 mM KCl, 1 mM EDTA, 1 mM dithiothreitol (DTT) supplemented with 0.04 % (w/v) n-Dodecyl-β-D-maltoside (DDM) was frozen in liquid nitrogen and stored at -80°C.

### Protein labelling

For SNAP labelling, 50 µg purified AHA2 were diluted with labelling buffer (50 mM MOPS-KOH, pH 7.0, 10 % (w/v) glycerol, 100 mM KCl, 1 mM DTT, 0.04 % (w/v) DDM) supplemented with 20 µM SNAP-Surface® Alexa Fluor® 647 (from a 1 mM stock in DMSO) and 1 g L^−1^ LUVs to a final concentration of 10 µM (final volume 40 µl) prior to incubation for 30 min on ice.

### Proteoliposome preparation

LUVs (4 - 5 mM) with 45 mM Octyl-β-D-Glucoside (OG) were incubated with end-over-end mixing for 5 min followed by addition of 25 µg of labelled AHA2 to a total volume of 200 µl. After incubating again, the sample was applied to a Sephadex G-50 fine spin column (3 ml matrix pre-equilibrated with reconstitution buffer). After 5 min incubation at room temperature (RT) the sample was centrifuged (8 min, RT, 180 x g, Eppendorf centrifuge 5810R) to separate proteoliposomes from the non-incorporated protein and detergent. Then, 100 mg Bio-Beads™ SM-2 (washed in methanol, then twice in ddH_2_O and stored in reconstitution buffer) were added to the flow-through and incubated at RT for 1 h with end-over-end mixing to remove residual detergent. The sample was separated from Bio-Beads™ by centrifugation (20 s, 200 x g, RT). Proteoliposomes were used within the next two days for microscopy and stored for characterization assays up to a week at 4°C.

### Proteoliposome immobilization

For TIRF microcopy, glass slides with 26 × 76 mm, #1.5 (Thermofisher) were cleaned by 2 × 15 min sonication in 2% Hellmanex™ III followed by methanol, respectively. Air dried glass slides were ozone cleaned in Novascan PSD Pro series Digital UV Ozone system (Novascan technologies, Boone, IA, USA) for 15 min and a bottomless 6 channel slide (Sticky-Slide Vl 0.4, Ibidi GmbH, Gräfelfing, Germany) was attached on the coverslip surface. Glass bottoms of channels were coated as described elsewhere (Thomsen et al., 2019). Briefly, 1 mg mL^−1^ biotinyl-PLL-PEG : PLL-PEG 1:100 was used for surface inactivation. After 20 min of incubation, unbound PEG was removed by 5 times washing with 60 µL reconstitution buffer followed by application of 0.025 g L^−1^ NeutrAvidin. After 20 min incubation and subsequent washing, proteoliposomes (2 - 5 μM, 1:1,000 dilution) were applied in channel and immobilization was allowed for 2 min. Afterwards, the channels were flushed 10 times with 60 µL reconstitution buffer to remove unbound liposomes. Controls without NeutrAvidin or without biotinyl-PEG DSPE show no fluorescence under our imaging settings, indicating an absence of non-specific binding of the liposomes.

### TIRF Microscopy

Images were obtained with an inverted total internal reflection fluorescence microscope (TIRF) model IX83 (Olympus, Hamburg, Germany). For acquisition, an oil-immersion objective (100×/1.5 numerical aperture (NA); Olympus) and an Orca-flash 4.0 sCMOS camera (Hamamatsu Photonics K.K., Hamamatsu, Japan) were used with an emission quad band filter cube (DAPI/FITC/CY3/CY5). The lipid bound NBD group and protein bound Alexa-647 were excited using lasers at 488 nm (143 µW) and 640 nm (104 µW), respectively. For all measurements, a penetration depth of 200 nm was applied to obtain images of 1,024 × 1,024 pixels, corresponding to 66.56 × 66.56 µm, with a dynamic range of 16-bit grayscale. The exposure time was set to 200 ms in both channels. For bleaching assays, three imaging rounds of the same 10 frames were taken using the multi-position tool with five images per frame in both channels. In between the runs, either reconstitution buffer (control), 10 mM dithionite (from 1 M stock in 500 mM Tris, pH 9.9) or 5 nM alamethicin (from 10 µM stock in ethanol with or without 10 mM dithionite) were flushed through the channel. After incubation of 2 min, the next round of images was taken. Z Drift compensation was used to keep the focal plane during the time course of the experiment. For photobleaching step analysis, five frame was imaged in the lipid channel 5 times and in the protein channel 1,000 times with 200 ms exposure time and 104 µW laser power.

### Image analysis

Images were converted to TIFF using FIJI (Schindelin et al., 2012) and analyzed with home written python algorithm, based on (Thomsen et al., 2019). First, fluorescent particles were detected and tracked over the time course of the experiment (up to 15 frames) for an x/y drift correction utilizing the trackpy module (Allan et al., 2021). The coordinates extracted from lipid channel were used to analyze the signals in the protein channel. The intensity of the detected particles and local background were extracted using the photutils and astropy modules (Astropy Collaboration et al., 2013, 2018) and background corrected. Since the method allows for parallel monitoring hundreds of individual proteoliposomes, a common drift was computed with trackpy, enabling the intensity extraction from the correct position, even if the signal of a spot decreased below the detection limit (Bohr et al., 2020, 2019; Thomsen et al., 2019).

Lipid and protein marker signals were pooled per set of five frames to a mean before/after#1/after#2 and filtered for data categorization. Only particles fulfilling certain criteria were taken for further analysis, namely LUV size (lipid marker signal), movement (standard deviation of signal in set of frames), tightness (lipid marker signal > 20%) after dithionite addition, complete bleaching after alamethicin addition in presence of dithionite (lipid marker signal < 10%) (Figure S1). LUVs were characterized based on the protein marker signal whether they are empty or protein containing. For orientation assay, only proteoliposomes above protein signal threshold of 250 A.U. were included. Photobleaching steps were analyzed with use of quickPBSA package (Hummert et al., 2021).

### Size distribution of liposomes

Sizes of proteoliposomes were calculated as described elsewhere (Hatzakis et al., 2009; Kunding et al., 2008; Lohr et al., 2009). Briefly, the fluorescence intensity of the lipid marker (*N*-NBD-DOPE) *I*_*NBD-DOPE*_ is proportional to the surface area *A* of the vesicle:

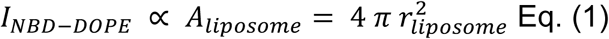

Assuming each liposome is a sphere with the radius *r*, equation 2 can be used to convert the intensity of the vesicle spot into the radius, using the calibration constant *k*_*cal*_ (Eq. 3).

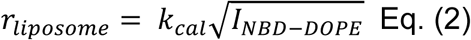

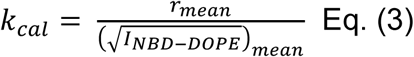

For each vesicle preparation, the actual mean radius *r*_*mean*_ of the preparation was determined by dynamic light scattering (DLS, Zetasizer, Malvern NanoZS, Worcestershire, UK) at 25 °C. The obtained autocorrelation function was converted into the size dependent intensity distribution with the Dispersion Technology Software, version 4.20 (Malvern Instruments Ltd., 2002).

### Poisson distribution calculation of protein

Based on the occurrence probability of differently sized liposomes, the theoretical occupancy of those vesicles with a specific protein copy number can be calculated assuming that the protein reconstitution efficiency is independent of the liposome size (Cliff et al., 2020). Briefly, the probability (Pr_n_) of a vesicle carrying specific copy number of proteins (n) is described by the Poisson distribution:

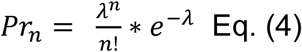

Here, *n* is the copy number of protein (occurrence of event) and *λ* is the Poisson-Parameter, which is equivalent to the expected value, in this specific case the average number of proteins per liposome:

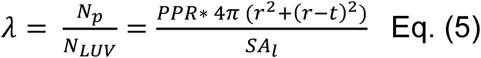

*N*_*p*_: number of proteins, *N*_*LUV*_: number of vesicles, *PPR*: protein per phospholipid ratio (*N*_*p*_*/N*_*l*_), *SA*_*l*_: lipid surface area, *r*: radius, *t*: bilayer thickness

The Poisson distribution can be further used to determine vesicle occupancy depending on the radius of a vesicle subpopulation *(Pr*_*n*_*(r))*:

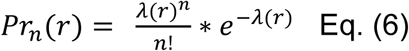

Therefore, the size distribution of the vesicles needs to be known to calculate the Poisson parameter *λ(r)* for the subpopulations:

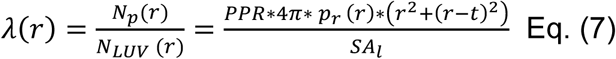

Finally, the obtained probability distribution Pr_n_(r) was multiplied by the corresponding probability of a radius to occur deriving the occurrence of a copy number at a given size.

### ATP-dependent proton transport assay

Acidification of vesicle lumens by AHA2 proton translocation was measured with ACMA fluorescence quenching assay (Dufour et al., 1982). Upon addition of MgSO_4_ (3 mM final concentration to 100 µM proteoliposomes in 20 mM MOPS-KOH, pH 7.0, 50 mM K_2_SO_4_, 3 mM ATP, 1 µM ACMA, and 62.5 nM valinomycin), ACMA fluorescence was quenched due to proton accumulation in the lumen. The H^+^-gradient was dissipated by the addition of 5 μM CCCP. Fluorescence signal was recorded over a period of 600 s at 480 nm (excitation 412 nm, slit width 2 nm, resolution 0.1 s) at 23 °C using a fluorometer (PTI-Quantamaster 800, Horiba, Benzheim, Germany). Fluorescence traces were normalized to the intensity measured directly after addition of MgSO_4_.

### Protein orientation bulk assay

Proteoliposomes were reconstituted the same way as described previously using unlabeled protein. Afterwards, the sample was split and incubated with the surface dye SNAP-Surface® Alexa Fluor® 488. After two hours, the membrane permeable dye SNAP-Cell® 647-SIR was added, and the samples were incubated again. As 100% value for comparison, one sample was incubated the whole time with the cell-permeable dye and another sample with the surface dye only, respectively. The samples were loaded on an SDS-PAGE and the gel was analyzed for in-gel fluorescence with the ChemiDoc MP imager (Bio-Rad, Hercules, CA, USA) using the Image Lab™ software and illumination at 460-490 nm and 520-545 nm with 530/28 nm and 695/55 nm emission filters, respectively, followed by colloidal Coomassie staining (Dyballa and Metzger, 2009) to verify that similar protein amounts were loaded. Signals were analyzed using Image Lab™ Software version 4.0 (Bio-Rad).

### Other Analytical Techniques

For protein quantification, 10 µl of proteoliposomes were loaded with BSA standard on a 10% SDS-PAGE followed by colloidal Coomassie staining. Phospholipid phosphorus was assayed after heat destruction in presence of perchloric acid as described previously (Chifflet et al., 1988). To verify detergent removal and lipid composition, vesicles were analyzed by thin-layer chromatography using chloroform:methanol:ammonium hydroxide (63:35:5, v/v/v) as solvent based on a modified protocol from (Eriks et al., 2003). Samples were applied without prior extraction by chloroform/methanol and detergent standards were applied to the same plate. For visualization, plates were stained with primuline (0.005% in acetone:water, 8:2; v/v) and imaged under long-wave UV light (ChemiDoc MP imager).

## Results and Discussion

### Design of the imaging assay for proteoliposomes

Our multi-parameter assay makes use of total internal reflection fluorescence (TIRF) microscopy to image, with high throughput, single proteoliposomes tethered to a solid support using particle tracking (shown schematically in Figures 1a and 2b) (Thomsen et al., 2019). TIRF microscopy was chosen because of reduced rate of photobleaching of the imaged fluorophores, lower background fluorescence, and higher signal-to-noise ratio compared with confocal microscopy. Tethering was accomplished with a biotin/neutravidin protocol, which maintains the native function and diffusivity of reconstituted transmembrane proteins (Mathiasen et al., 2014), and the spherical morphology (Bendix et al., 2009) and low passive ion permeability (Li et al., 2015; Guha et al., 2021) of the vesicles. Purified plant H^+^-ATPase AHA2 was used as a model transporter. It functions as an ATP-driven membrane intrinsic protein that plays a key role in the physiology of plants by controlling essential functions such as nutrient uptake and intracellular pH regulation (Palmgren, 2001; Serrano et al., 1986). Proteoliposomes were formed by insertion of Alexa-647-labelled H^+^-ATPase AHA2 into liposomes containing trace quantities of the fluorescent marker lipid *N*-NBD-DOPE and a biotinylated anchor lipid (Figure S2a). Thin layer chromatography (TLC) confirmed efficient removal of detergent (Figure S2b). The resulting liposomes displayed ATP-dependent H^+^-transport activity (Figure S2c).

**Fig. 1:**
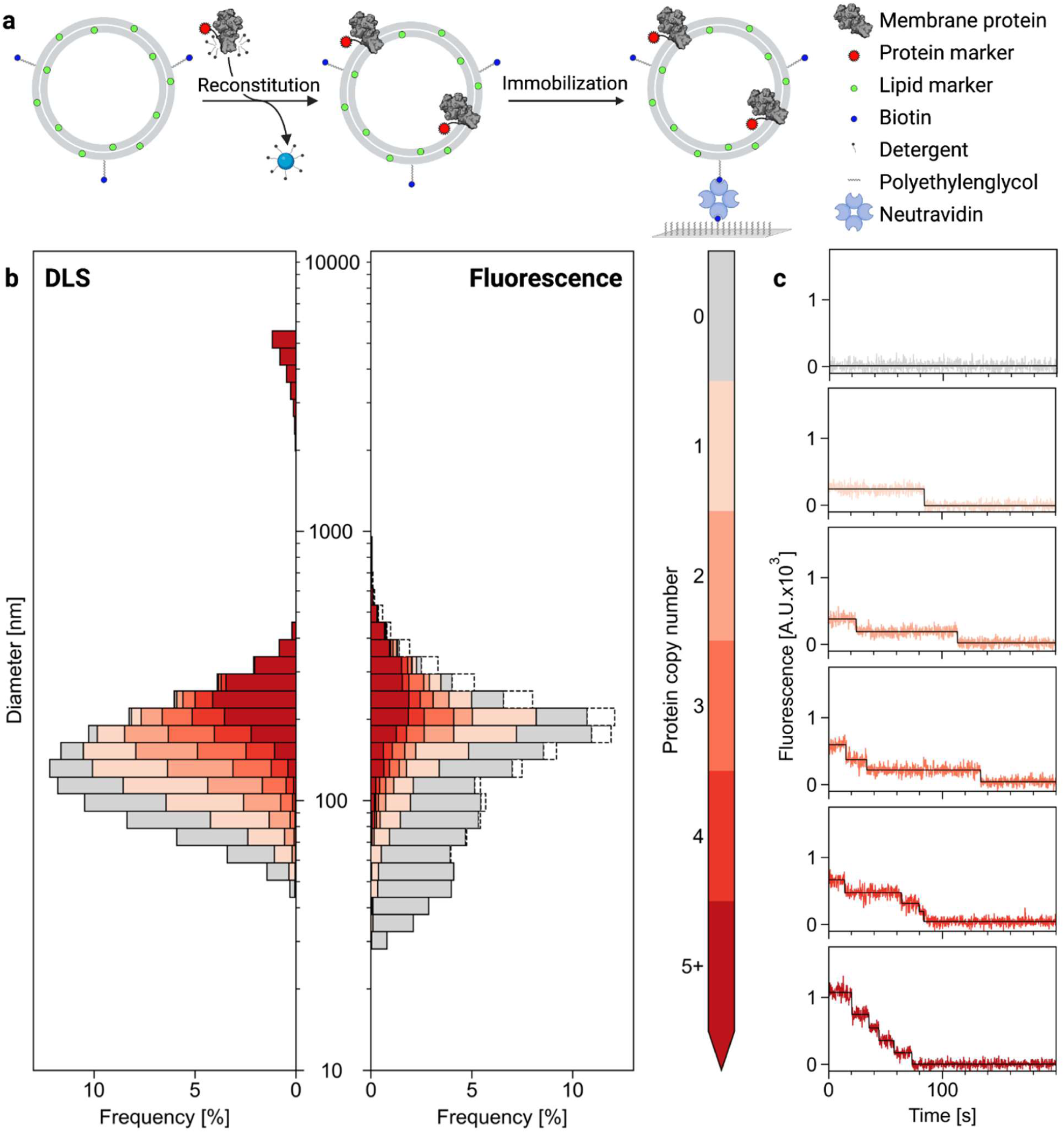
Generation and analysis of proteoliposomes. (a) Schematic illustration of the reconstitution procedure. Preformed liposomes containing trace amounts of Biotinyl-PEG-DSPE and the fluorescent marker lipid *N*-NBD-DOPE are detergent-destabilized and mixed with detergent-solubilized Alexa-647-labelled H^+^-ATPase AHA2. Detergent removal results in the formation of sealed proteoliposomes, which are immobilized for imaging via TIRF microscopy. (b) Representative histograms displaying the size distribution of liposomal preparations after reconstitution, as determined from dynamic light scattering (DLS, left) or single vesicle analysis (Fluorescence, right). The expected protein distribution based on bulk protein and lipid determination (left) is compared with measured protein content based on photobleaching analysis of 3,287 vesicles (right). Dashed lines in the right panel indicate additional detected vesicles of specific size where the bleaching trace could not be fitted successfully. (c) Exemplary bleaching traces for each vesicle category (1-5+ proteins; black lines represent the fitted average fluorescence intensity). Control studies confirmed that vesicles prepared without protein did not display fluorescence in the protein channel and allowed the determination of a protein signal threshold (see Figure S4a).

**Fig. 2.**
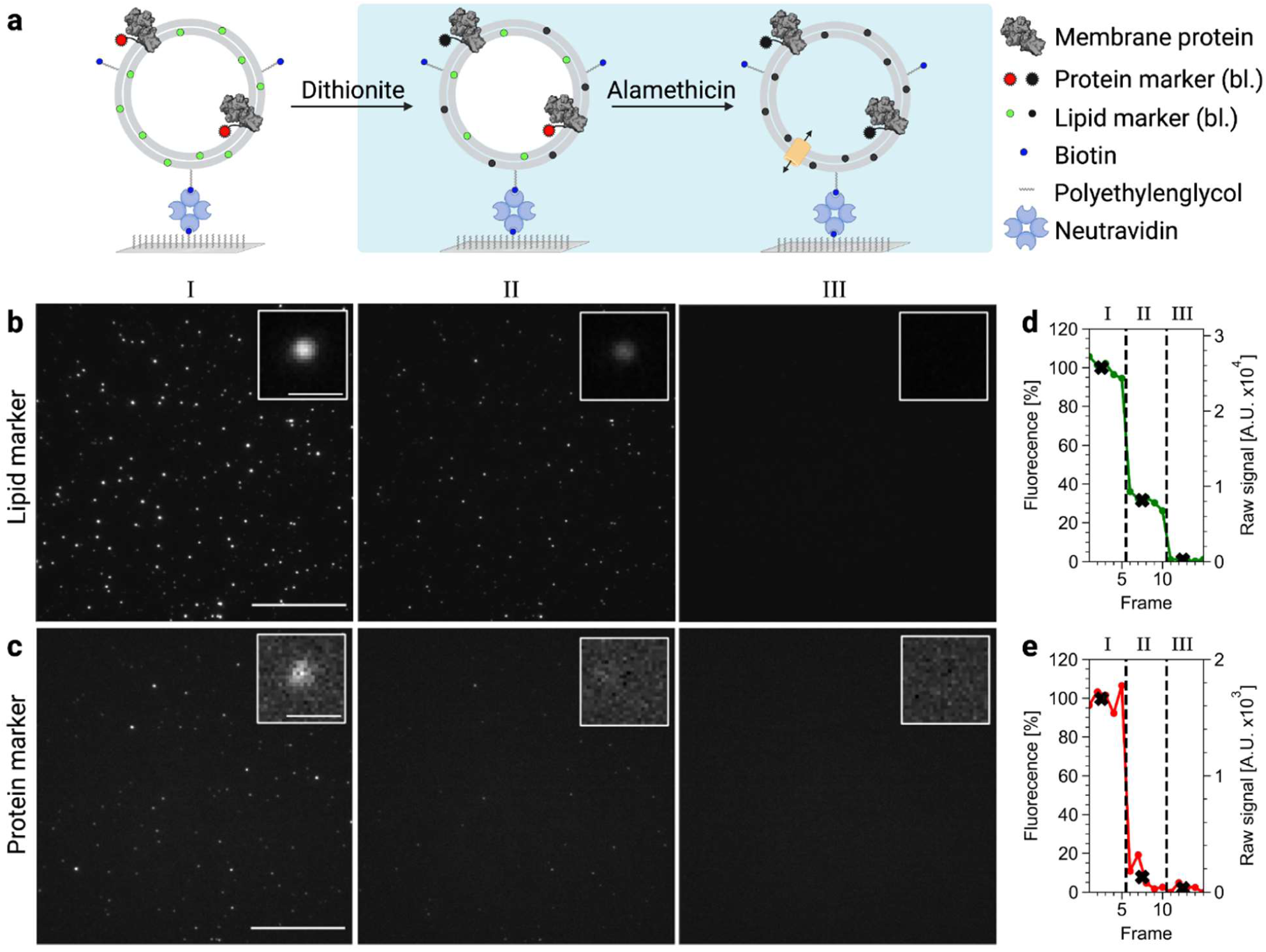
Single vesicle fluorescence-bleaching assay for multi-parameter analysis. (a) Schematic illustration of the bleaching assay. Immobilized liposomes prepared with fluorescently tagged membrane proteins (Alexa647-AHA2) and lipid markers (*N*-NBD-DOPE) are exposed to dithionite to irreversible quench the fluorescence signal of outward facing labels. Addition of the pore forming peptide alamethicin allows dithionite to reach all fluorophores and serves as control for complete bleaching (bl.). (b, c) Exemplary TIRF images of immobilized vesicles in the lipid channel (b, NBD) and protein channel (c, Alexa647) before (I), after dithionite addition (II) and subsequent addition of alamethicin (III). Scale bar, 20 µm. *Inset:* exemplary proteoliposome. Scale bar, 1 µm. (d, e) Fluorescence intensity traces of the exemplary proteoliposome in the lipid channel (d, green trace) and protein channel (e, red trace), respectively. Five images were recorded per each step of the experiment. The black crosses indicate the mean values for each step of the experiment. For the selected vesicle, dithionite addition leads to total loss of the protein signal although a signal remains in the green channel, indicating tight vesicles with outward orientated protein.

DLS characterization based on scattering intensity showed liposomes with intensity-averaged diameters in the range of 51 to 460 nm with a small number (2.9 ± 1.4 %, n=3) of larger aggregates (>1,000 nm) (Figure 1b). We used the size distribution determined by DLS in combination with protein determination via quantitative SDS-PAGE and lipid determination via phosphate assay (protein/lipid ratio 0.86 mg mmol^-1^) to calculate a theoretical protein distribution among the vesicle population (Figure 1b, left panel), based on the Poisson equation (Cliff et al., 2020; Stockbridge, 2021). We note that this calculation assumes a size independent reconstitution efficiency of protein in liposomes.

We next immobilized the vesicles onto passivated glass slides (Figure 1a) and recorded fluorescence images of the lipid and protein channels. The images were analyzed by custom-written software to detect individual liposomes and their protein content, while excluding aggregates (Figure S1). Using the mean liposomal diameter determined by DLS, the lipid signal distribution of the immobilized liposomes was converted into a vesicle size distribution (Hatzakis et al., 2009), (Figure 1b, left panel). Compared to the analysis in bulk, we observed for the immobilized liposomes an increase in the number of smaller vesicles (cf. Figures 1b, left and right panel). This enrichment of small vesicles, noted in previous studies (Olsson et al., 2015; Hatzakis et al., 2009), is likely because they diffuse more quickly and thereby immobilize faster as the vesicle suspension flows over the slide. By analyzing the number of steps in the fluorescence bleaching traces of the spots from TIRF microscopy images in the protein channel (Figure 1c), we could deduce the number of proteins in each vesicle and obtain the experimental protein distribution (Figure 1b, right panel). For vesicle with up to four proteins, the photobleaching traces allowed resolving the exact copy numbers per single liposome (Figure 1c). The experimental distribution showed a higher number of protein-free vesicles in comparison to the calculated protein distribution based on bulk analysis, likely because of the higher number of smaller vesicles and the exclusion of aggregates in the single vesicle analysis that are included in the bulk assay analysis. Notably, we observed protein-free vesicles of both small and large sizes, consistent with previous reports based on ensemble measurements of the presence of a subpopulation of vesicles that is resistant to protein reconstitution (Niu et al., 2009; Ploier et al., 2016; Lee et al., 2009; Stockbridge, 2021). Likewise, the number of proteins per vesicle differed substantially, highlighting the heterogeneity of the reconstituted preparation, in line with previous reports (Pick et al., 2018; Cliff et al., 2020). These differences between theoretical predictions and experimental data clearly illustrate the limitation of ensemble measurements on proteoliposomes regarding proper interpretation of results.

### Single vesicle-based bleaching assay

We next made use of the membrane-impermeant dianion dithionite to bleach fluorescently labeled lipids and labeled protein reconstituted into liposomes, thereby providing information on the intactness and lamellarity of the vesicles, and the transbilayer orientation of the protein. On adding dithionite, we observed irreversible bleaching of the outward facing NBD fluorophores, while inner-leaflet fluorophores remained protected in many vesicles (Figure 2a and b, middle panels). Subsequent addition of the pore-forming antibiotic peptide alamethicin enabled dithionite to enter the vesicles, thereby bleaching luminally-oriented fluorophores and eliminating all fluorescence in the image (Figures 2a and b, right panels). Quantification of the lipid fluorescence of an exemplary liposome is depicted in Figure 2d (further examples are shown in Figure S3). As can be readily observed (Figure 2d), addition of dithionite caused a sharp drop in NBD fluorescence to ∼40% of its starting value. Addition of alamethicin caused a further sharp drop, resulting in elimination of fluorescence. The NBD fluorophore is susceptible to photobleaching, a problem that we could not eliminate despite imaging under low photobleaching conditions via TIRF. However, the photobleaching was reproducible under our conditions (fluorescence loss per set of 5 frames was 11 ± 5%; mean ± SD, n = 1,997) and could therefore be readily incorporated into subsequent analyses as a quantitative correction (see later). We conclude that the vesicle highlighted in the boxed inset is sealed to dithionite permeation and unilamellar. More extensive analyses of the vesicle population are presented in the next section.

Parallel imaging of the immobilized vesicles in the protein channel (Alexa-647; Figure 2c) allowed identification of protein-containing vesicles and determination of the accessibility of the protein marker. For the same exemplary liposome analyzed above and shown to be sealed and unilamellar (boxed in Figure 2b and c), complete loss of protein fluorescence was observed upon dithionite addition (Figure 2e), indicating an exclusive orientation of the reconstituted protein with the fluorescently labeled SNAP-tagged N-terminus accessible to the extravesicular space. We note that addition of dithionite or alamethicin did not affect vesicle immobilization on the slide (Figures S4b and c).

### Multi-parameter analysis of proteoliposomes

Using the single liposome approach, we imaged thousands of vesicles before and after addition of dithionite and first analyzed the accessibility of the lipid marker (Figure 3a and Figure S5). We observed two populations of vesicles upon dithionite addition. A minor population (Population I, comprising ∼28% of all vesicles) displayed a total loss of the lipid signal on dithionite addition, indicating leakiness. However, most of the vesicles (∼72%), Population II, displayed a drop in NBD fluorescence to ∼30% of the starting value. Correcting for ∼10% photobleaching as noted above, this result indicates that ∼40% of the lipid marker is in the inner leaflet of this vesicle population, which is somewhat lower than the estimated 48% for vesicles of diameter >100 nm. We thus conclude that most of the imaged vesicles, comprising population II, are sealed to dithionite and unilamellar.

**Figure 3.**
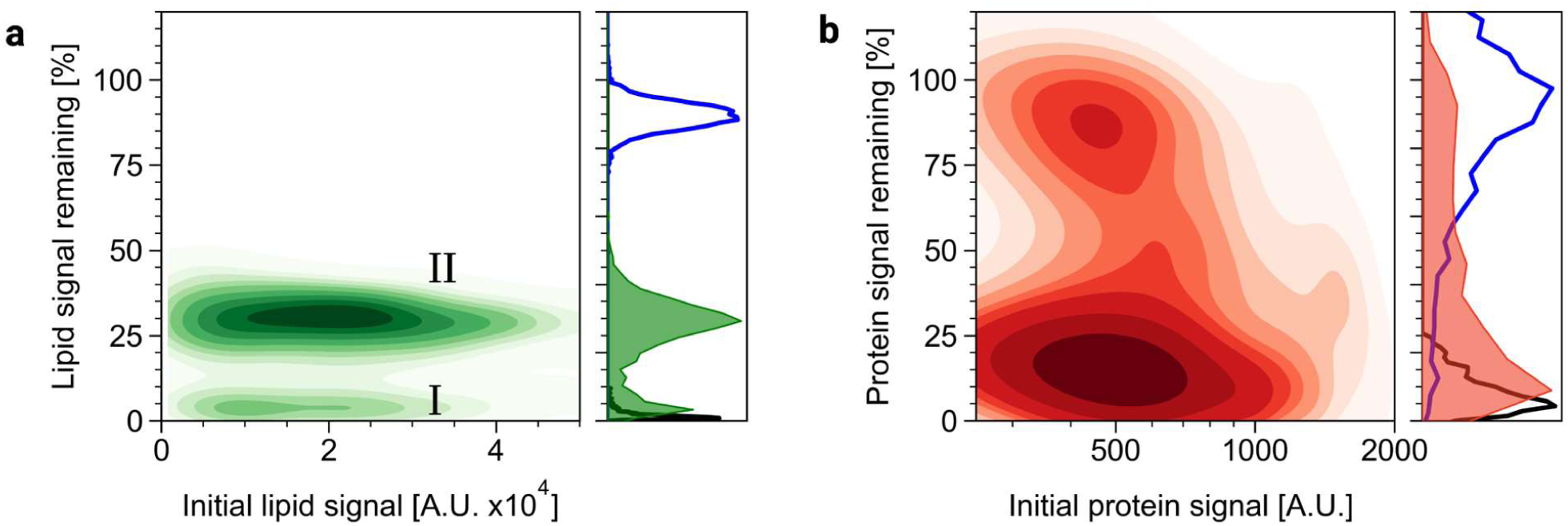
Multi-parameter analysis of proteoliposomes. Immobilized proteoliposomes prepared with fluorescently tagged membrane proteins (Alexa647-AHA2) and lipid markers (*N*-NBD-DOPE) were imaged by TIRF microscopy. (a) Kernel density estimation plot of the remaining lipid signal vs. initial lipid signal upon addition of dithionite. Two separate subpopulations (I and II) are visible. (B) Subpopulation II was analyzed for the protein signal before and after dithionite addition; shown is the kernel density estimation plot. Distribution plots are shown on the right of each panel; for comparison line plots of control measurements are included (blue, buffer control; black, alamethicin in presence of dithionite, see Figure S5). Data are based on bleaching analysis of at least 1,300 protein-containing liposomes (protein signal > 250 A.U.).

Population II of tight vesicles was further analyzed for protein marker accessibility. Two subpopulations were observed preferably within the class of proteoliposomes with a low amount of reconstituted protein (Figure 3b). The major subpopulation (51%) showed a strong loss of the protein signal (to less than 30% of the original) upon dithionite addition and thus represents vesicles with proteins displaying their N-terminal SNAP tag outwards. The minor subpopulation (23%) displayed a stable protein signal (>70% of the initial intensity) after dithionite addition that overlapped with the buffer-treated sample (Figure 3b and Figure S5), representing inward-facing proteins. For vesicles containing a high amount of reconstituted protein, dithionite addition resulted in 0-100% loss of fluorescence, reflecting a mix of inside and outside oriented proteins in these vesicles. A side-by-side comparison with an ensemble sidedness assay showed 88% outside orientation of the reconstituted protein (Figure S2d), as compared to the averaged estimation (67%) from the single vesicle analysis. Such asymmetric reconstitution is occasionally observed for membrane proteins with a large cytoplasmic part and is depending on the liposomal lipid composition (Hickey and Buhr, 2011; Marek et al., 2011) and purification conditions (Niu et al., 2002).

For further analysis, we plotted the signals for the lipid marker vs. protein marker remaining after dithionite addition for all imaged vesicles considering their different protein content (Figure 4a). This allows extraction of valuable additional information regarding the different proteoliposome subpopulations. First, vesicles that lose both lipid and protein signals must be leaky. This leaky vesicle subpopulation comprises both low and high protein-containing liposomes and thus is an intrinsic result of the reconstitution protocol, possibly including subsequent immobilization on the microscope slide. Second, by plotting the signals for remaining lipid signal upon addition of dithionite vs. square root of initial lipid signal for all imaged vesicles considering their different protein content it becomes evident that the leaky subpopulation comprised both small and large vesicles without any preference for vesicle subpopulation of small diameter, excluding curvature dependency (Figure 4b). Third, it is also evident that the preparation contains few multilamellar proteoliposomes, in particular vesicles with larger diameter, as indicated by their remaining lipid signal intensities >40% upon dithionite addition.

**Figure 4.**
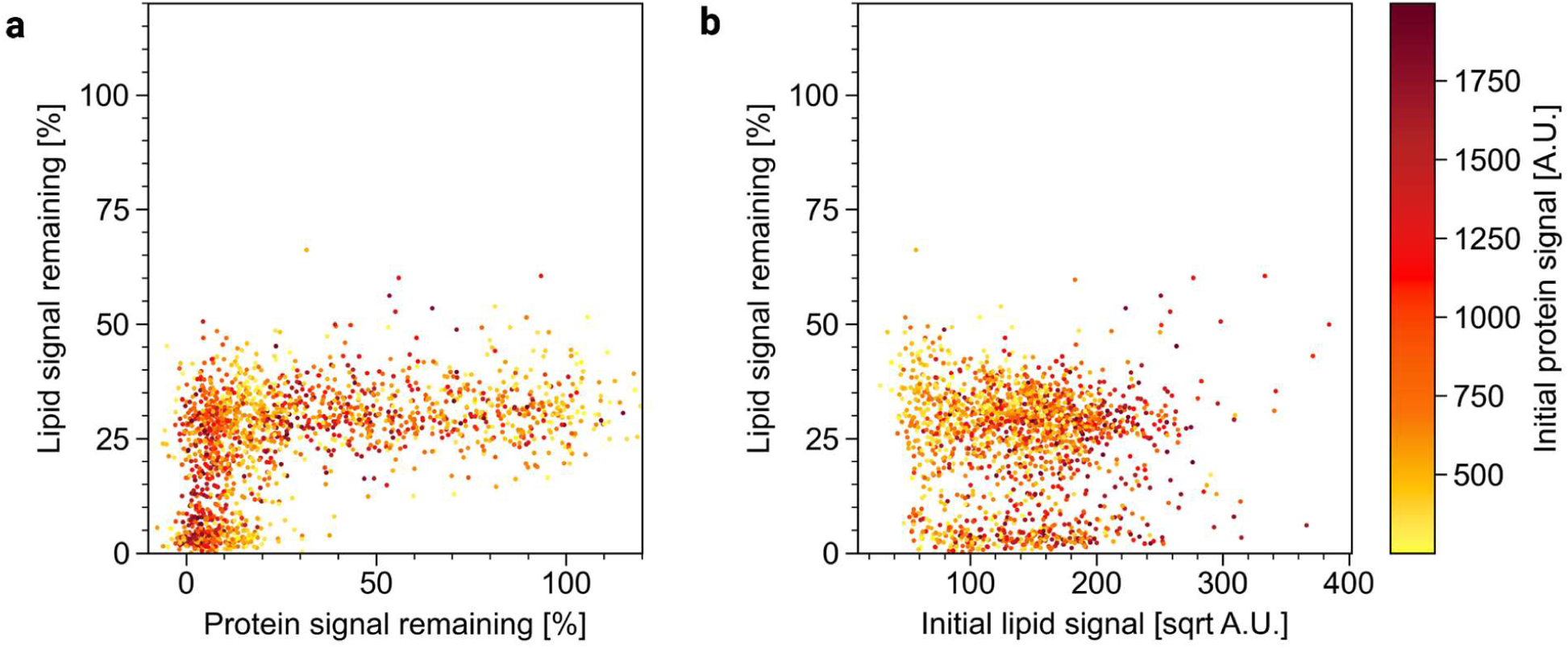
Characterization of proteoliposome subpopulations by multi-parameter analysis. Immobilized liposomes were prepared with fluorescently tagged membrane proteins (Alexa647-AHA2) and lipid markers (*N*-NBD-DOPE). (a) Protein-containing liposomes (protein signal > 250 A.U.) are plotted for their lipid signal remaining vs. protein signal remaining after dithionite addition. The subpopulation dropping to zero shows neither remaining lipid nor protein signals, indicating leakiness. (b) Protein-containing liposomes (protein signal > 250 A.U.) are plotted for their remaining lipid signal vs. square root of initial lipid signal upon addition dithionite. The leaky subpopulation comprises both small and large vesicles, excluding curvature effects. The colormap indicates the initial protein signal.

## Conclusion

We describe a two-color TIRF microscopy-based bleaching protocol to analyze individual vesicles reconstituted with fluorescent lipids and membrane proteins. Using custom software, we could analyze the images to yield multiple physical parameters of individual proteoliposomes, including unilamellarity, protein content and protein orientation. We could define a sub-population of vesicles that was leaky, thereby excluding them from downstream analyses. Likewise, aggregates and non-incorporated protein were also analytically excluded, thereby avoiding the need for additional purification procedure, e.g. liposome floatation, prior to vesicle capture. With the large range of fluorophores commercially available, we anticipate that our approach can be expanded from dual to multi-color setup to enable quantitative, parallel analyses including determination of membrane compositional heterogeneity and physical properties. The versatile strategy presented here is a powerful tool for optimization of reconstitution conditions and should benefit further investigation of many types of membrane proteins, including analysis of thus far under-studied lipid flippases at the single vesicle level (Thomsen et al., 2019; Shukla and Baumgart, 2021; López-Marqués et al., 2020).

## Supporting Information

Description of images analysis; ensemble characterization of the liposomal preparations; exemplary time traces of single vesicles; control studies on immobilized vesicles; multi-parameter assay controls on immobilized proteoliposomes.

## Acknowledgements

This work was funded by grants from the German Research Foundation (INST 213/886-1 FUGG, INST 213/985-1 FUGG; GU 1133/11-1) and DAAD (57386621) to TGP, and the National Institutes of Health (Grant EY027969) to AKM. SV and LCP gratefully acknowledge funding from Studienstiftung des deutschen Volkes. NSH acknowledges funding from Novo Nordisk Foundation (grant numbers NNF16OC0021948 and NNF14CC0001) and the Villum Foundation (BioNEC grant 18333). The authors thank Anne-Mette Bjerg Petersen for excellent technical assistance, Bo Justesen for providing the AHA2 expression plasmid, and Eckhard Hofmann for access to DLS. Open access funding enabled and organized by Project DEAL. Figures were created with BioRender.com.

## Author Contributions

TGP and AKM conceived and designed the project. TGP and NSH supervised the research. SV and LCP conducted all the experiments, developed analysis algorithms and wrote the Python scripts. NSH and SB contributed to TIRF imaging and data analysis. SV and LCP performed data analysis with assistance from SB. SV, LCP, AKM and TGP wrote the manuscript. All authors discussed the results and commented on the manuscript.

