## Supporting information for "A single vesicle fluorescence-bleaching assay for multi-parameter analysis of proteoliposomes by total internal reflection fluorescence microscopy"

This file includes all Supporting Information: Description of images analysis; ensemble characterization of the liposomal preparations; exemplary time traces of single vesicles; control studies on immobilized vesicles; multi-parameter assay controls on immobilized proteoliposomes.

**Particles** (7,919)

- Lipid signal detectable in all 5 initial images
- Separation = 30 pixel

→ **Liposomes** (4,132)

Initial lipid signal is:

- > 650 A.U.
- Fluctuation < 7.5 % of average

Remaining lipid signal after dithionite and alamethicin addition < 10%

↓ **Proteoliposomes** (1,803)

Initial protein signal is:

- > 250 and < 2,000 A.U.
- Fluctuation < 50% of average

Remaining protein signal after dithionite and alamethicin addition < 25%

← **Tight Proteoliposomes** (1,302)

Lipid remaining signal after SDT addition > 20%

**Figure S1. Description of image analysis.** Images were processed and analyzed in four steps using in-house developed routines in python. For every filtering step, total amount of particles is noted. In the first step, all fluorescent particles with a separation of 30 pixels were considered if the lipid signal (NBD) was detectable in all five consecutive initial images. In the second step, fluorescent particles were identified as liposomes if both the averaged NBD fluorescence intensity was > 650 A.U. with a standard deviation < 7.5% and less than 10% of the initial NBD fluorescence intensity was left on addition of alamethicin in presence of dithionite. In the third step, liposomes were further classified as proteoliposomes if the protein signal (Alexa647) displayed an averaged fluorescence intensity between > 250 and < 2,000 A.U. with a maximum standard deviation of 50%. Additionally, the protein signal after dithionite and subsequent alamethicin addition had to be < 25% of initial protein signal. In the last step, proteoliposomes were classified as tight if the lipid signal on dithionite addition remained > 20% of the initial signal.

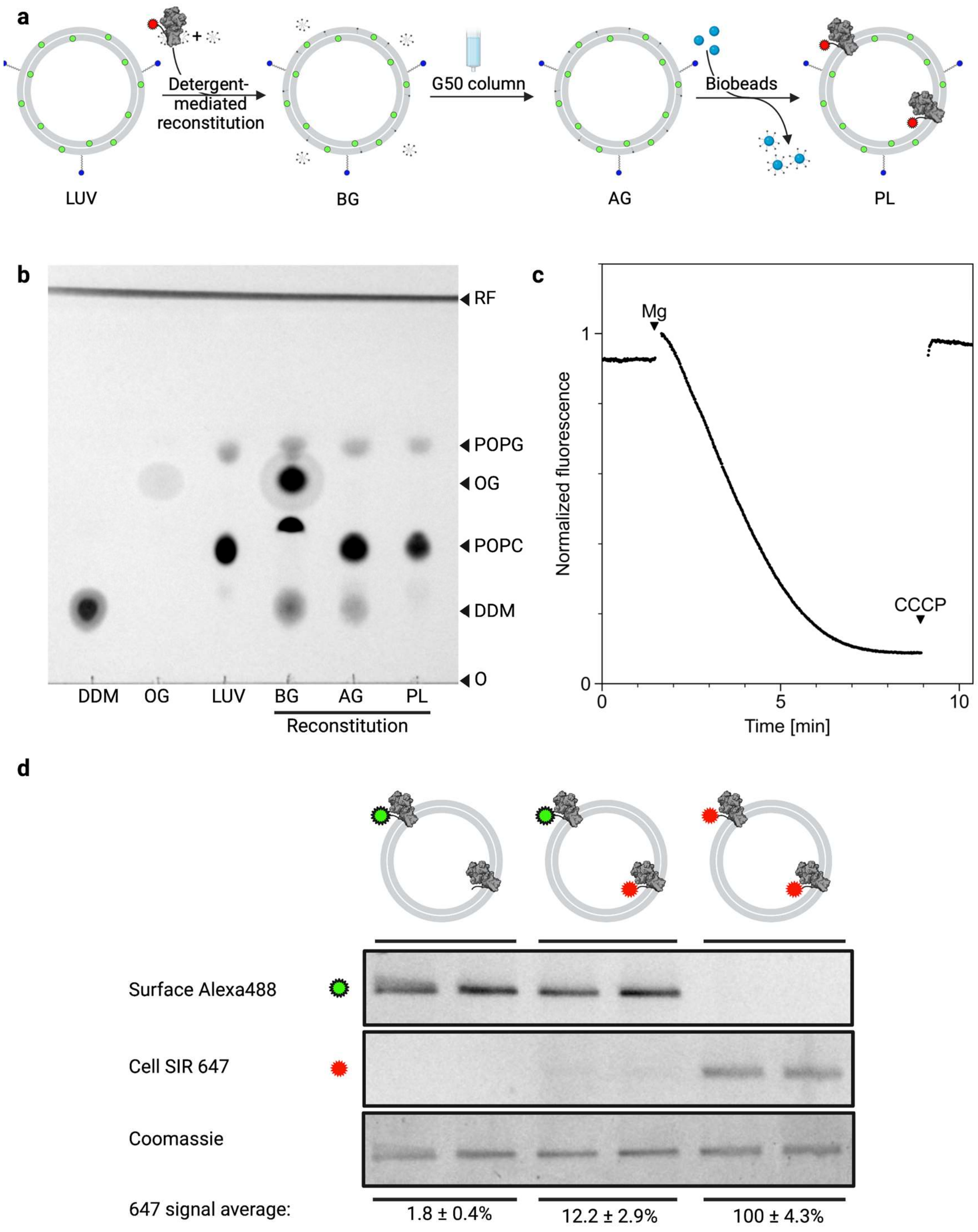


**Figure S2. Ensemble characterization of the liposomal preparations.** (a) Schematic illustration of the reconstitution procedure. Preformed large unilamellar liposomes (LUV) containing trace amounts of Biotin-PEG-DSPE and the fluorescent marker lipid *N*-NBD-DOPE are detergent-destabilized and mixed with detergent-solubilized membrane proteins ($H^+$-ATPase AHA2) fluorescently tagged with Alexa-647 (BG). Subsequent removal of the detergent by Sephadex G-50 gel filtration (AG) followed by Bio-Bead treatment result in the formation of sealed proteoliposomes (PL). (b) Thin layer chromatography analysis of samples (LUV, BG, AG, PL)

from different steps during proteoliposome reconstitution revealed efficient detergent removal. Samples were resolved onto a silica plate along with detergent standards (50 nmol; DDM, n-dodecyl-β-D-maltoside; OG, n-octyl-β-D-Glucoside) and imaged under UV light after staining with primuline. POPC, 1-palmitoyl-2-oleoyl-*sn*-glycero-3-phosphocholine; POPG, 1-palmitoyl-2-oleoyl-*sn*-glycero-3-phospho-(1'-rac-glycerol); RF, running front; O, origin. Note that the presence of significant amounts of detergents affects separation (lane BG). (c) $H^+$ pumping in reconstituted proteoliposome vesicles initiated by addition of $MgSO_4$ to reconstituted proteoliposomes in buffer containing the fluorophore ACMA, valinomycin, and ATP. The decrease in ACMA fluorescence reflecting proton accumulation into the lumen of the vesicles was abolished by addition of the protonophore CCCP. (d) Ensemble protein sidedness assay. Unlabelled SNAP-AHA2 was reconstituted into liposomes and subsequently labelled separately or sequentially with membrane impermeable and membrane permeable SNAP dyes (SNAP-Surface® Alexa Fluor® 488 and SNAP-Cell® 647-SIR, respectively). Samples were analyzed by SDS-PAGE followed by in-gel fluorescence determination followed by Coomassie staining to verify equal amount of protein in all samples. Estimation of membrane orientations of AHA2 revealed that about 88% of the protein faced outward with their cytoplasmic portion.

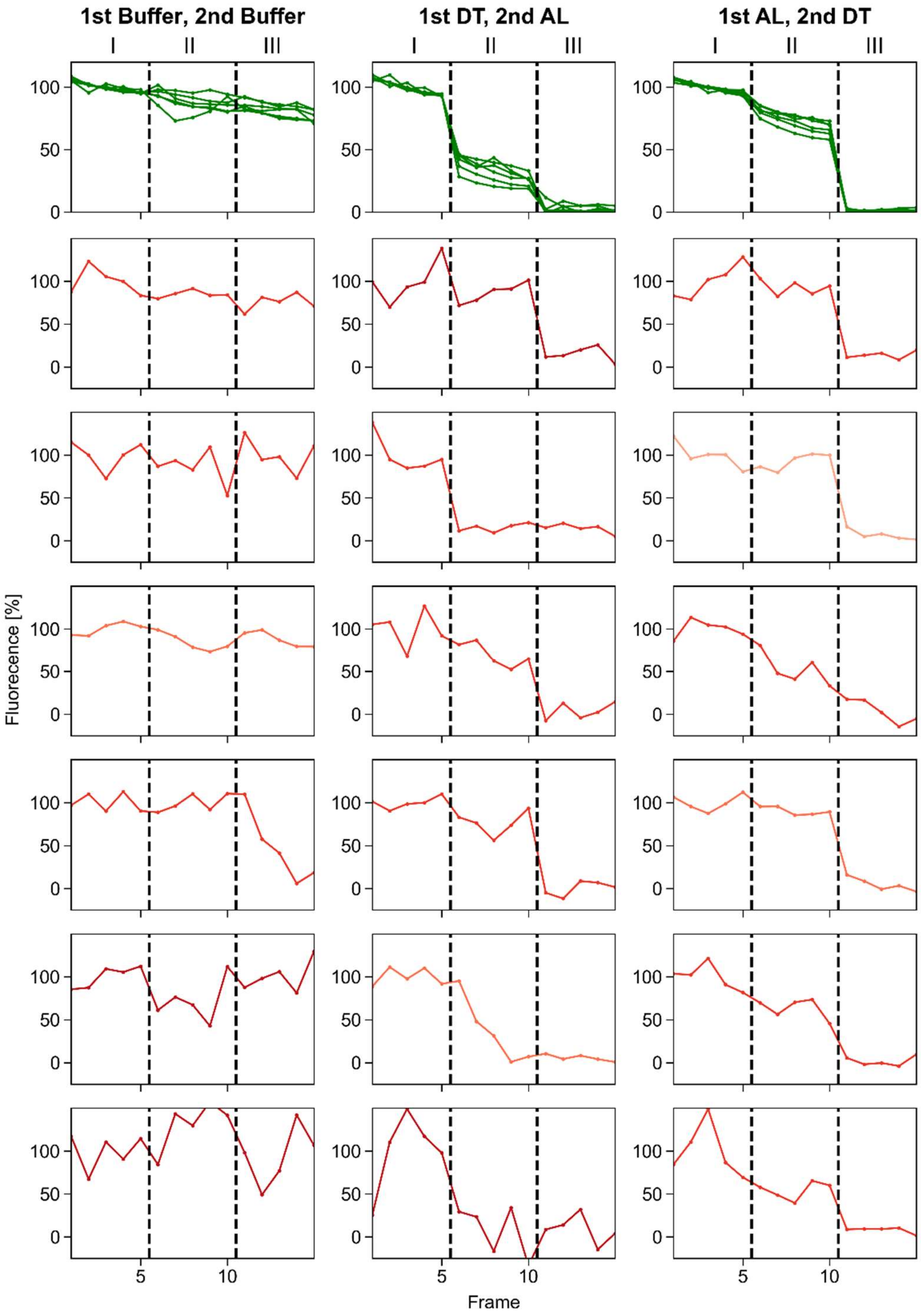


**Supplementary Figure S3: Exemplary time traces of single vesicles.** Immobilized liposomes prepared with fluorescently tagged membrane proteins (Alexa647-AHA2) and lipid markers (*N*-NBD-DOPE) were imaged by TIRF microscopy. Exemplary fluorescence intensity traces of six single vesicles in the lipid channel (NBD, green traces) and protein channel (Alexa647, red, the darker the higher the initial signal) are shown per condition. The dotted lines mark the periods before (I), after 1st addition (II) and after 2nd addition (third column) as indicated. Five images were recorded per each step of the experiment. DT, dithionite; AL, alamethicin.

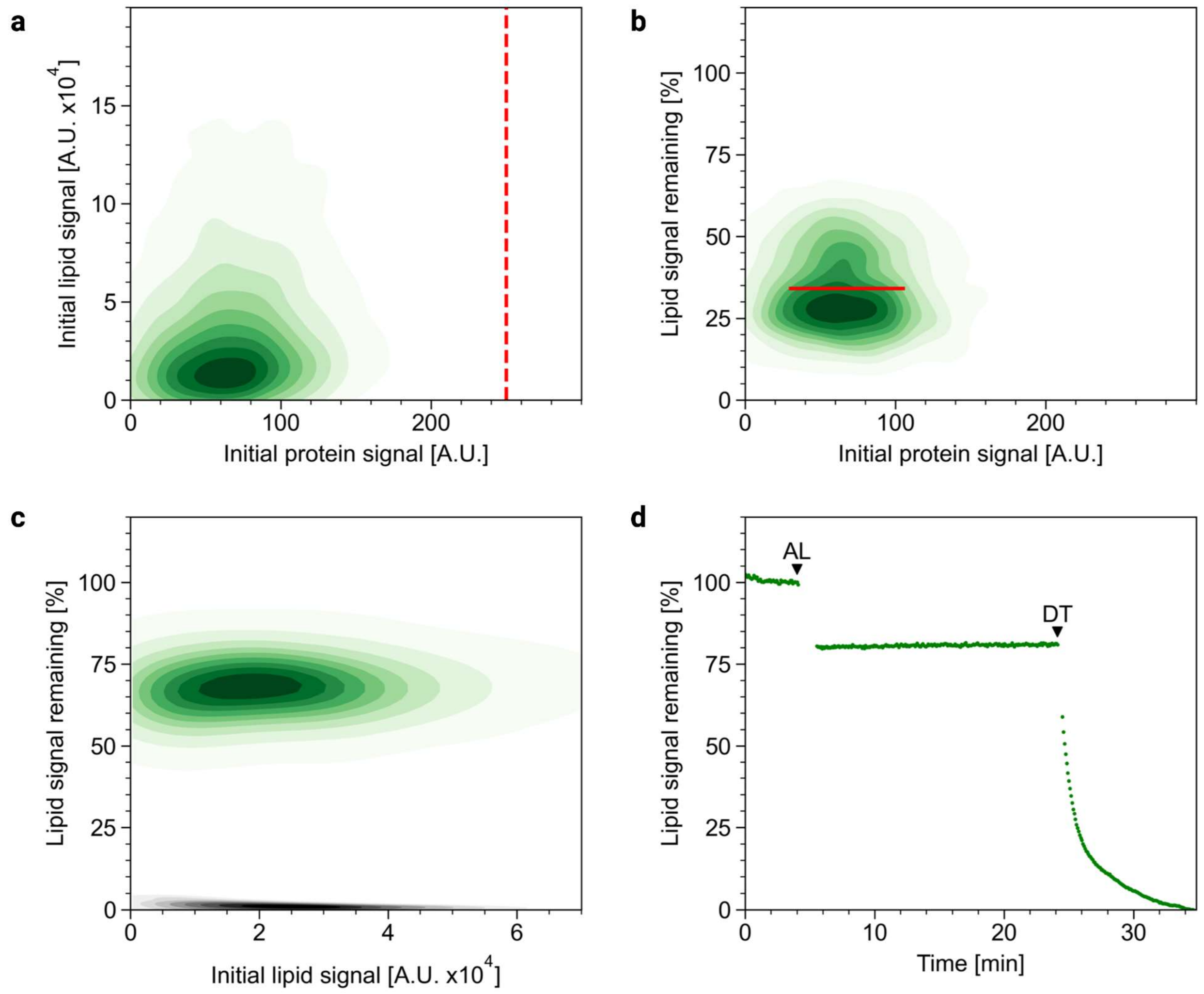


**Figure S4. Control studies on immobilized vesicles.** (a, b) Immobilized LUVs prepared with fluorescently tagged lipid markers (*N*-NBD-DOPE) by manual extrusion were imaged by TIRF microscopy. (a) Kernel density estimation plot of the initial NBD fluorescence detected in the lipid channel vs. protein channel showing no correlation between both fluorescence signals, thus demonstrating the absence of bleed-through lipid fluorescence into the protein channel. The vertical dashed line corresponds to 250 A.U. representing the threshold value above background intensity levels in the protein channel. (b) Kernel density estimation plot of the remaining NBD fluorescence vs. the initial signal in the protein channel upon addition of dithionite. All vesicles remained visible ruling out that dithionite is affecting vesicle immobilization. Red line indicates mean lipid fluorescence upon dithionite addition (34.2 %). (c, d) Liposomes were prepared with fluorescently tagged membrane proteins (Alexa647-AHA2) and lipid markers (*N*-NBD-DOPE). (c) Analysis by TIRF microscopy. Kernel density estimation plot of the remaining NBD fluorescence upon alamethicin addition vs. the initial signal in the lipid channel (green, average 69.4 %) followed by dithionite addition (black). (d) Ensemble measurements utilizing a plate reader CLARIOstar®. The NBD fluorescence was excited at 460 nm and the emission recorded at 536 nm. The arrowheads indicate addition of alamethicin (AL, 5 nM final concentration) followed by dithionite

(DT, 10 mM final concentration); note that the dilution effect was negligible. In both measurements (panels c and d), addition of alamethicin permeabilizes the liposomes and causes a decrease in NBD fluorescence by affecting the photophysical properties of the lipid-linked NBD group at the liposomal membrane. However, the remaining signal demonstrates that alamethicin is not affecting vesicle immobilization. Subsequent addition of dithionite resulted in complete elimination of fluorescence, demonstrating that the dithionite concentration is sufficient to reduce completely the NBD groups after 2 min of incubation.

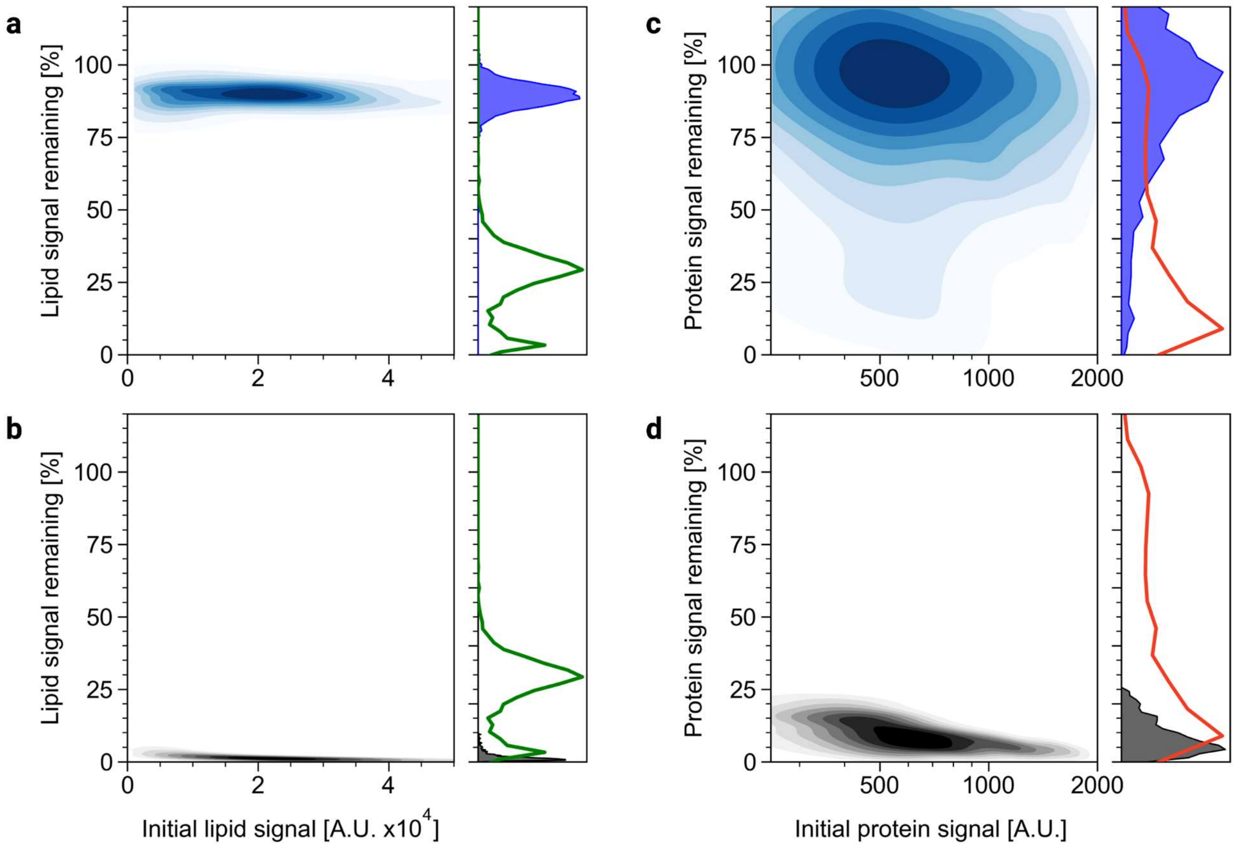


**Figure S5. Multi-parameter assay controls on immobilized proteoliposomes**. Immobilized proteoliposomes prepared with fluorescently tagged membrane proteins (Alexa647-AHA2) and lipid markers (N-NBD-DOPE) were imaged by TIRF microscopy. Kernel density estimation plots of the remaining lipid signal vs. initial lipid signal (a,b) and the remaining protein signal vs. initial protein signal (c, d) upon addition of buffer (a, c) and alamethicin in presence of dithionite (b, d). Distribution plots are shown on the right of each panel; for comparison line plots of the remaining signals upon dithionite addition are included (green, lipid signal; red, protein signal, see Figure 3). Data are based on bleaching analysis of at least 1,300 protein-containing liposomes (protein signal > 250 A.U.).
